# Predicting emergent phenotypes from single cell populations using CELLECTION

**DOI:** 10.1101/2025.09.02.673886

**Authors:** Hongru Hu, Siddhant Sanghi, Gerald Quon

## Abstract

Biological systems exhibit emergent phenotypes that arise from the collective behavior of individual components, such as whole-organ functions that arise from the coordinated activity of its individual cells, or organism-level phenotypes that result from the functional interplay of collections of genes in the genome. We present CELLECTION, a deep learning framework that learns to associate subgroups of instances with different emergent phenotypes. We show CELLECTION enables interpretable predictions for heterogeneous tasks, including disease classification, identification of disease-associated cell subtypes, alignment of developmental stages between human model systems, and even predicting relative hand-wing indices across the avian lineage. CELLECTION therefore provides a scalable and flexible framework for identifying key cellular or genetic signatures underlying complex traits in development, disease, and evolution.

## Main

Biological traits arise from a cascade of interactions that spans multiple layers of biological systems. Variants in DNA, regulatory programs encoded in chromatin, and external cues are integrated through the circuitry of single cells before emerging as sample-level phenotypes. Because each biological layer can amplify, buffer or redirect the signal from the previous layer, the rules governing how individual biological components give rise to emergent phenotypes is complex^1–4^. Unraveling these rules that connect phenotypes to fundamental units of biological systems is a central goal of modern genomics.

For example, consider Alzheimer’s disease (AD). AD arises from a complex interplay of gene regulatory alterations across multiple brain cell types, which together drive the neurodegenerative cascade^5–8^. In neurons, dysregulation of synaptic and metabolic gene expression impairs plasticity and energy homeostasis^9–11^, while in microglia, disease-associated transcriptional states are marked by upregulation of *TREM2*, *TYROBP*, *APOE*, and interferon-responsive genes that promote a pro-inflammatory environment and impaired clearance of amyloid-β plaques^12–15^. Astrocytes show aberrant activation and altered cholesterol metabolism gene expression, contributing to neuronal stress and blood-brain barrier dysfunction^16–19^. Meanwhile, oligodendrocyte lineage cells demonstrate downregulation of myelination and lipid biosynthesis programs, possibly affecting white matter integrity and neural connectivity^20–23^. These coordinated yet cell type-specific gene regulatory changes converge to amplify neuronal dysfunction, synapse loss, and neuroinflammation, thereby driving AD progression.

Most single cell genomics technologies available to identify cell type-specific changes associated with pathologies such as AD are applied to each cell type individually, and do not identify coordinated changes across cell types^24,25^. For example, after collecting snRNA-seq data from a case-control cohort, one strategy is to cluster cells and use cell markers to identify cell types, compute cell-type proportions per individual sample, then identify cell types that change significantly in proportion between cases and controls. This cell type proportion strategy has two major limitations: (i) it assumes that all cells of the same type, and all samples within a phenotypic class are biological replicates of the same homogeneous state, and (ii) it relies on accurate cell clustering and cell type annotation, which are often inconsistent across phenotypic classes and require substantial reference alignment and manual curation. Another strategy is the identification of cell type-specific differentially expressed genes (DEGs), which again requires that cells first be separated into discrete cell types and groups before identification of DEGs. This approach discards information about cell type heterogeneity and can yield misleading results if the specific cell states that drive differences between cases and controls are only a small subpopulation of a given cell type.

Importantly, cell type proportion and DEG analysis are both unable to identify coordinated changes amongst multiple cell types that, only when occurring together, drive case-control status. To bridge this gap, we introduce CELLECTION, a deep learning framework that models each biological sample as an unordered set of individual cells, and identifies specific combinations of cell states that, when present in the same sample, and predictive of sample-level phenotypes. We first demonstrate CELLECTION achieves state-of-the-art performance at using scRNA-seq data to predict binary patient case-control phenotypes in COVID-19^26^, lupus^27^ and cardiomyopathy^28^ data, as well as continuous phenotype prediction tasks such as COVID-19 severity^29,30^ and developmental age^31,32^. We then apply CELLECTION to identify universal signatures of brain development that are consistent between brain organoids and fetal brain samples, even when the underlying scRNA-seq data from each type of sample shows poor overlap in cell states. Second, we perform a deep analysis of Alzheimer’s disease case-control snRNA-seq data^25^ that significantly refines a previously-identified disease-associated microglia signature associated with AD pathology. Finally, we show CELLECTION has application areas well beyond single cell biology, by demonstrating it can predict hand-wing index variation across hundreds of avian species, based only on gene content of their genomes. These results demonstrate CELLECTION’s ability to extract robust, biologically meaningful cellular programs that generalize across datasets, systems, and phenotypic contexts.

## Results

### CELLECTION predicts emergent phenotypes from collections of molecular instances

CELLECTION is a deep learning framework based on point clouds^33^ that models biological samples as unordered collections of molecular instances, and learns to predict sample-level phenotypes using only molecular features of the instances. In the context of this paper, we focus on two applications: predicting patient-level case-control phenotypes based on collections of cells transcriptionally sequenced from each individual, and predicting organism-level phenotypes of a species, given the entire collection of genes encoded in its genome. For brevity, in the remainder of this text we will mainly describe CELLECTION from the view of predicting patient-level phenotypes from collections of single cells, though our framework logic equally applies to organism phenotype prediction from genomes.

The intuition of CELLECTION is best understood in the context of case-control prediction, such as predicting Alzheimer’s disease status as presented in the latter portion of this paper. In a collection of sequenced nuclei from a single patient, each sequenced nucleus represents the molecular state of a single cell and its corresponding cell type in high-dimensional gene expression space. Each cell type corresponds to a specific region of this gene expression space, and within a given region for a cell type, some subregions may correspond to healthy cell states, while others correspond to disease cell states. During training to predict Alzheimer’s disease status, CELLECTION learns to identify which combinations of regions of gene expression space are associated with Alzheimer’s patients but not healthy controls, and makes corresponding predictions of sample-level phenotypes. From the viewpoint of a Uniform Manifold Approximation and Projection (UMAP)^34^ visualization of single nuclei data, CELLECTION intuitively identifies regions of UMAPs that, when present together in an individual, are strongly associated with the Alzheimer’s phenotype. Importantly, by not collapsing the snRNA-seq instance-level measurements to pseudo-bulk averages at the sample- or cell type level, CELLECTION processes the full set of cells directly, identifying phenotype-relevant regions of the cellular landscape without relying on manual clustering or annotation. As we demonstrate later in the manuscript, this allows CELLECTION to learn novel sub-cell types discriminative of cases and controls without prior cell type annotation.

Formally, CELLECTION makes predictions at the sample-level by transforming individual instances (cells) into low-dimensional embeddings before attention-based^35^ aggregation and prediction (**Figure.1**). Every cell in a sample is first independently passed through a shared-weight multilayer perceptron (MLP) that applies a nonlinear transformation of the input features (**Figure.1a**). Alternatively, instead of optimizing its own instance encoder, CELLECTION can apply a pretrained cell encoder such as sciLaMA^36^ to take advantage of transfer learning from large single cell atlases. Because the same network is applied to each cell, the model respects permutation invariance and can generalize across samples with differing numbers of cells. Afterwards, CELLECTION applies a sequence of Feature Transformation Blocks, each primarily consisting of a Transformation Network (T-Net) module^37^, which learns an affine transformation matrix that aligns the input point cloud to a standard pose across samples in the training dataset. This module can stabilize training and improve generalization by reducing sample-specific distortions in cellular feature space due to sample-level covariates.

**Figure. 1.**
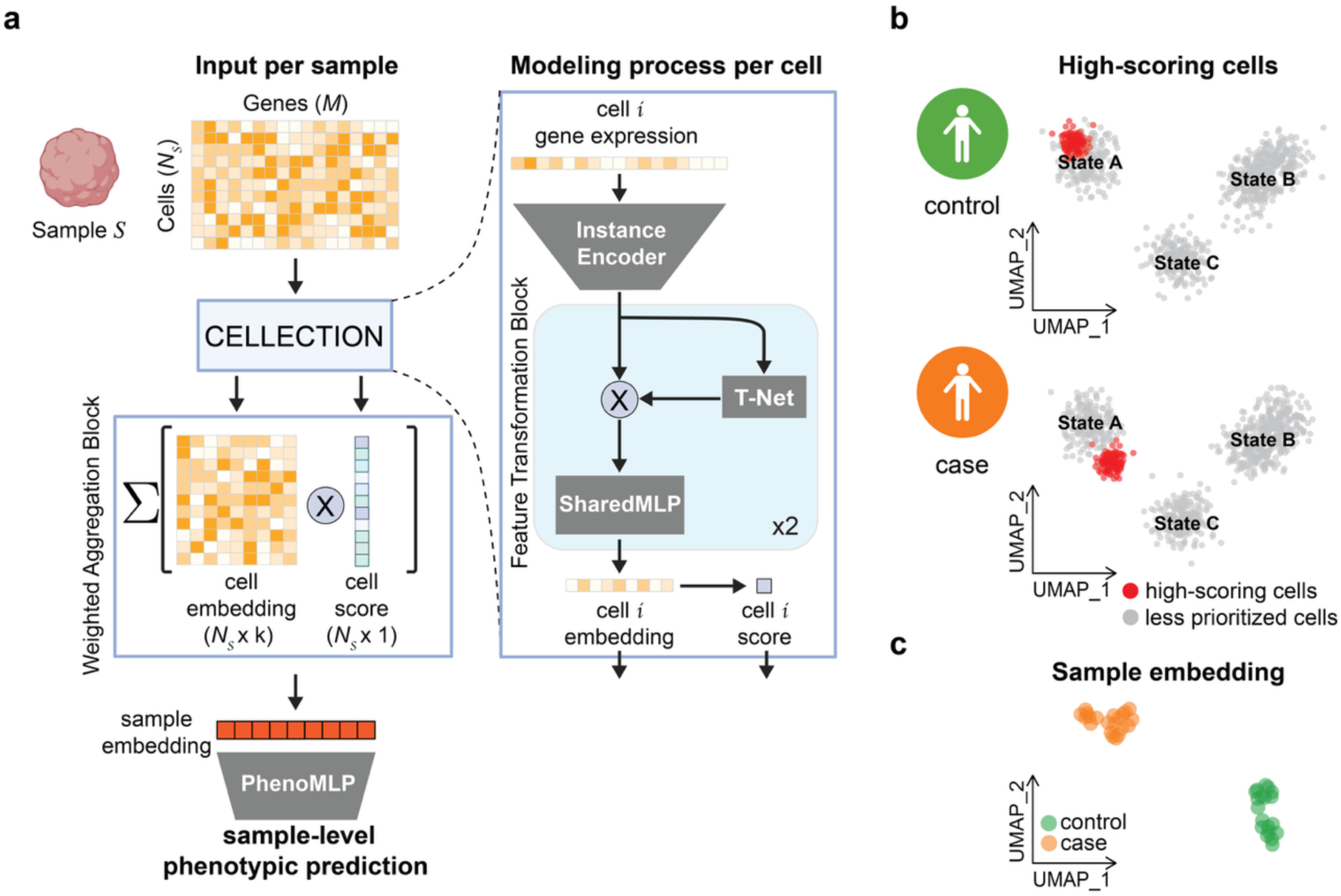
Overview of the CELLECTION framework. (**a**) Schematic of the CELLECTION model architecture and data flow. Each biological sample *S* is represented as a variable-sized set of single-cell transcriptomes (*N*_;_cells × *M* genes). An instance encoder and sequence of Feature Transformation Blocks processes each cell *i* into a cell-level embedding, before a Weighted Aggregation Block computes an associated importance score based on an attention mechanism and combines the cell embeddings via a weighted aggregation mechanism to derive a sample-level embedding. This sample embedding is used for phenotype prediction through a multilayer perceptron (PhenoMLP). (**b**) An example UMAP visualization of all cells used as input to CELLECTION across all input samples, where each cell is labeled by its importance score assigned during training. In our visualizations, cells within the top 10% of scores are highlighted in red, to illustrate which cells are most prioritized for phenotype prediction. There are different sets of cells prioritized for each phenotype being predicted (cases and controls separately in this example). (**c**) An example UMAP visualization of sample embeddings learned by CELLECTION, with each point representing an individual sample (collection of cells) and colored by control-case status.

After processing individual cells via the Feature Transformation Blocks, CELLECTION aggregates them within a Weighted Aggregation Block into a single sample-level representation using learned cell-level attention scores (**Figure.1b**). Instead of summarizing cells through naive averaging or max-pooling, the model computes a learned importance score for each cell, which reflects its relevance to the phenotype prediction task. These scores are used to compute a weighted average of the cell local features, allowing the model to focus on key subpopulations that contribute to the sample-level outcome. This attention-based aggregation is central to CELLECTION’s interpretability, as it enables identification of phenotype-relevant cells without any prior knowledge of cell types or marker genes. The resulting sample-level embedding is then passed through a prediction head implemented as a fully connected network for classification or regression, depending on the specific task. Importantly, the number of cells can vary across input samples, and predictions are invariant to the order of the instances in the data. The entire model is trained end-to-end in a weakly-supervised manner based on the phenotype signal at the sample level.

### CELLECTION accurately predicts disease status from scRNA-seq data

We first evaluated the performance of CELLECTION across five classification tasks derived from three population-scale scRNA-seq datasets. These tasks involve prediction of donor-level disease state and characteristics such as ethnicity, allowing us to assess the model’s robustness across biological contexts and outcome types. The datasets include a COVID-19 study from Ziegler et al.^26^, a lupus cohort from Perez et al.^27^, and a cardiomyopathy cell atlas from Chaffin et al.^28^. From these datasets, we defined five distinct prediction tasks: COVID-19 status (case vs. control), lupus nephritis status (systemic lupus erythematosus (SLE) case vs. control), donor race (Asian vs. White, within the lupus cohort), and cardiomyopathy status (non-failing (NF) hearts, dilated cardiomyopathy (DCM), and hypertrophic cardiomyopathy (HCM)), using either donor-level samples (“Cardio”) or biosample-level samples (“Cardio (biosample-level)”), the latter reflecting the availability of multiple biopsies for the same individual donor. To contextualize CELLECTION’s performance, we compared its prediction performance, interpretability and generalizability against several approaches used for sample-level phenotype prediction from single-cell data. These include cell type proportion (CT proportion) analysis, which computes the relative abundance of annotated cell types per sample; pseudo-bulk averaging (pseudobulk), which aggregates all single-cell profiles within a sample into a single expression vector; and cell-type-specific pseudo-bulk averaging, which averages expression within each annotated cell type per sample. We also included DeepSet^38^, a permutation-invariant neural network that aggregates instance-level features using a sum pooling operation; CloudPred^39^, a model that assumes each cell is sampled from a mixture of Gaussians, assigns cells to latent subpopulations, and predicts sample-level phenotypes based on the estimated prevalence of these subpopulations; and ProtCell4P^40^, a prototype-based neural network^41–43^ that learns representative cell prototypes in a latent space and classifies sample-level transcriptomic collections based on their distances to these prototypes.

Across datasets and prediction tasks, we found CELLECTION consistently achieved state of the art performance and more stable calibration in both Area Under the Receiver Operating Characteristic curve (AUROC) and Macro F1 metrics (**Figure.2a, b**). More specifically, CELLECTION outperformed other methods by an average of 12% in AUROC (p-value = 2.43e-29, paired Wilcoxon test), and an average of 46% in Macro F1 (p-value = 5.56e-39, paired Wilcoxon test). CELLECTION was significantly more predictive than cell type proportion analysis (mean AUROC increase of 20%, mean Macro F1 increase of 108%, p-value = 4.12e-30 and 2.52e-40, respectively, paired Wilcoxon test), indicating that CELLECTION captures significantly more information than just differences in cell type proportions across phenotypes. CloudPred exhibited instability across datasets, with large performance variation depending on cohort structure and class balance. ProtCell4P, a prototype-based method that relies on predefined cell type annotations and the number of prototypes, consistently underperformed relative to CELLECTION, although its performance increases as the number of prototypes increased. Importantly, CELLECTION achieves the best performance without requiring any annotation supervision or prior knowledge of cell types. DeepSet also performed well (ranking second on average based on Macro F1 score) but lacked interpretability due to its architectural design, which treats all instances equally through sum pooling and does not expose instance-level contributions. This limitation is critical in biological contexts, where identifying the specific cells or subpopulations driving a phenotype is essential for uncovering underlying mechanisms and generating biological insights. Generally, cell type proportion, pseudobulk, and cell-type-specific pseudobulk methods often achieved relatively high AUROC scores but performed poorly on Macro F1 metrics. This discrepancy stems from class imbalance, where AUROC remains robust to skewed class distributions, while Macro F1 penalizes models that fail to recover minority classes. As a result, these methods tend to overfit to the majority class, leading to inflated AUROC values that mask poor classification performance across all classes. In contrast, methods such as CELLECTION, DeepSet, and ProtCell4P exhibited more stable performance across both metrics, suggesting better generalization and more balanced phenotype prediction.

**Figure. 2.**
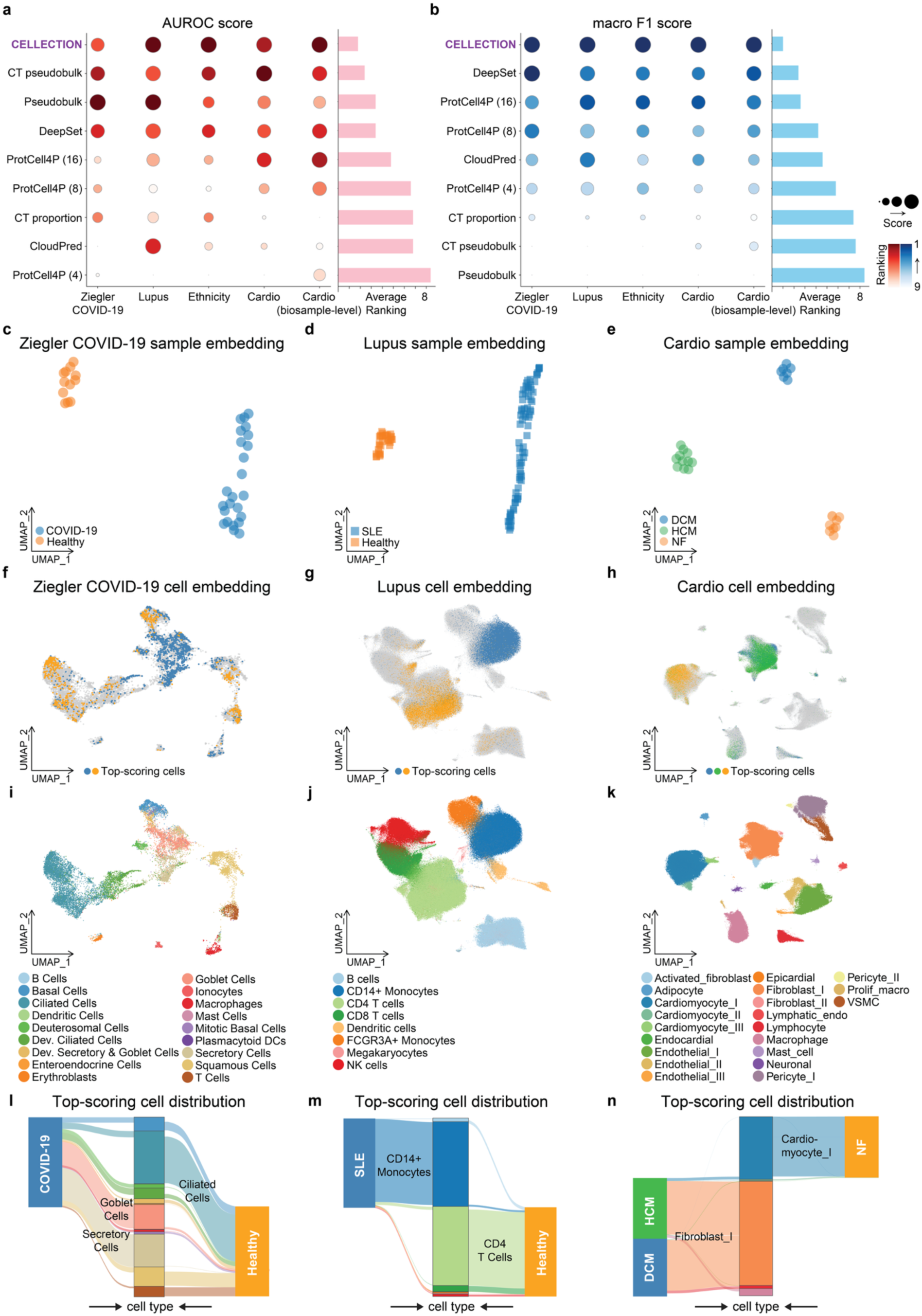
Benchmarks for sample-level phenotypic prediction and cell-level prioritization. (**a**) Benchmarking of phenotype prediction performance across multiple datasets or tasks (columns) and methods (rows), evaluated using AUROC on held-out test data. Dot size reflects scaled AUROC score (larger indicates better performance), while dot color represents ranking (darker indicates better rank). The bar plot on the right summarizes each method’s average rank across all datasets, where lower rank is better. (**b**) Same as (**a**), but evaluated using macro F1 score. (**c-e**) Sample-level embeddings learned by CELLECTION for the Ziegler COVID-19, Lupus, and Cardiac datasets, colored by annotated phenotypes. (**f-h**) Cell-level embeddings for the same datasets, with the top 10% high-scoring cells colored by their associated phenotype. (**i-k**) Cell-level embeddings colored by cell type annotations from the original studies. (**l-n**) Sankey plots showing enrichment of specific cell types among the top-scoring cells for each phenotype in the Ziegler COVID-19^26^, Lupus^27^, and Cardiac^28^ datasets, respectively.

We next explored the cell states that CELLECTION identified through attention in order to achieve its high classification accuracy, and determined the extent to which there existed heterogeneity in attended cells within each class label (cases or controls). CELLECTION estimates a sample-level embedding that summarizes the entire population of cells that were input for a single sample, as well as a per-cell score that indicates the importance of each cell to the phenotype prediction. At the sample level for these benchmark datasets, CELLECTION’s learned embeddings exhibited clear structure, with distinct separation between phenotype classes when projected into a low-dimensional global feature space (**Figure.2c-e**). This clear separation of phenotype classes is consistent with CELLECTION’s high prediction accuracy.

In the Ziegler COVID-19 cohort, CELLECTION prioritized goblet and secretory cells as key contributors to disease classification (**Figure.2f, i, l** and **Figure.S1a, d**). These goblet and secretory cells exhibit high expression levels of Angiotensin-Converting Enzyme 2 (ACE2)^26,44^, the key cellular entry receptor for SARS-CoV-2 that facilitates viral binding, attachment, and subsequent internalization into host cells. Note that CELLECTION is significantly more predictive than cell type proportion predictors for COVID-19 classification (mean AUROC increase of 11%, mean Macro F1 increase of 37%, p-value = 0.046 and 1.05e-5, respectively, paired Wilcoxon test), indicating that CELLECTION is identifying specific cell signatures within these two cell populations. CELLECTION’s *de novo* prioritization of goblet and secretory cells as key contributors to COVID-19 classification therefore suggests that the model effectively recovers known cellular targets of viral entry.

In the lupus dataset, CELLECTION consistently upweighted CD14+ monocytes cells as predictive of disease status (**Figure.2g, j, m** and **Figure.S1b, e**). This aligns with previous findings that type I interferon-stimulated genes (ISGs) are elevated in monocytes from SLE patients^27,45^, reflecting an activated interferon response.

In the cardiac dataset, both dilated cardiomyopathy (DCM) and hypertrophic cardiomyopathy (HCM) samples were driven by signals from fibroblast populations, while non-failing (NF) control samples were associated with cardiomyocyte signatures (**Figure.2h, k, n** and **Figure.S1c, f**). These results are concordant with the original study^28^, which reported that DCM and HCM hearts showed a significant reduction in cardiomyocyte abundance and increases in activated fibroblasts compared with NF hearts. Particularly, DCM hearts showed increased abundance in fibroblasts, macrophages, and lymphocytes, which is also well captured by our cell prioritization result (**Figure.S1f**). Such shifts in cellular programs were effectively recapitulated by CELLECTION’s attention-based cell prioritization.

Taken together, these results show that CELLECTION not only achieves state of the art prediction performance across multiple classification settings, but also offers interpretable, attention-guided insights into the cellular drivers of sample-level phenotypes.

### Sample level embeddings capture continuous disease progression

Many phenotypes of interest in biology and medicine lie along a continuum, including examples such as disease severity, developmental stage, and aging. In such settings, classification loss functions such as those used in the previous section treat all misclassifications equally, instead of considering the relative position of the predicted and measured outcome along the continuum. To model such ordered outcomes, we extended CELLECTION with ordinal regression objectives, enabling it to learn from rank-based phenotype labels and generate interpretable predictions even in the absence of sharp class boundaries.

We first applied the ordinal regression approach to predict COVID-19 severity using single cell profiles from the large-scale Haniffa group dataset^30^ (**Figure.3a, b**), which includes more individual samples (n=120) than the aforementioned Ziegler data^26^. In the Haniffa dataset, donors were annotated along an eight-point severity scale ranging from healthy to death. CELLECTION, trained with an ordinal regression objective, accurately recovered the underlying progression, achieving strong correlation between predicted and true severity levels in the test set (Spearman = 0.71) (**Figure.3c**). The resulting sample-level embeddings formed a continuous manifold where samples aligned smoothly along the disease progression axis (**Figure.3d**), and exhibited better topological continuity in the sample-level embedding space than the classification model (**Figure.S2**). Notably, the model’s attention scores highlighted gradual shifts towards monocytes and plasmablast cell states from B cells as severity increased (**Figure.3e** and **Figure.S3a, b**), revealing a dynamic immunological response aligned with clinical outcome trajectories.

**Figure. 3.**
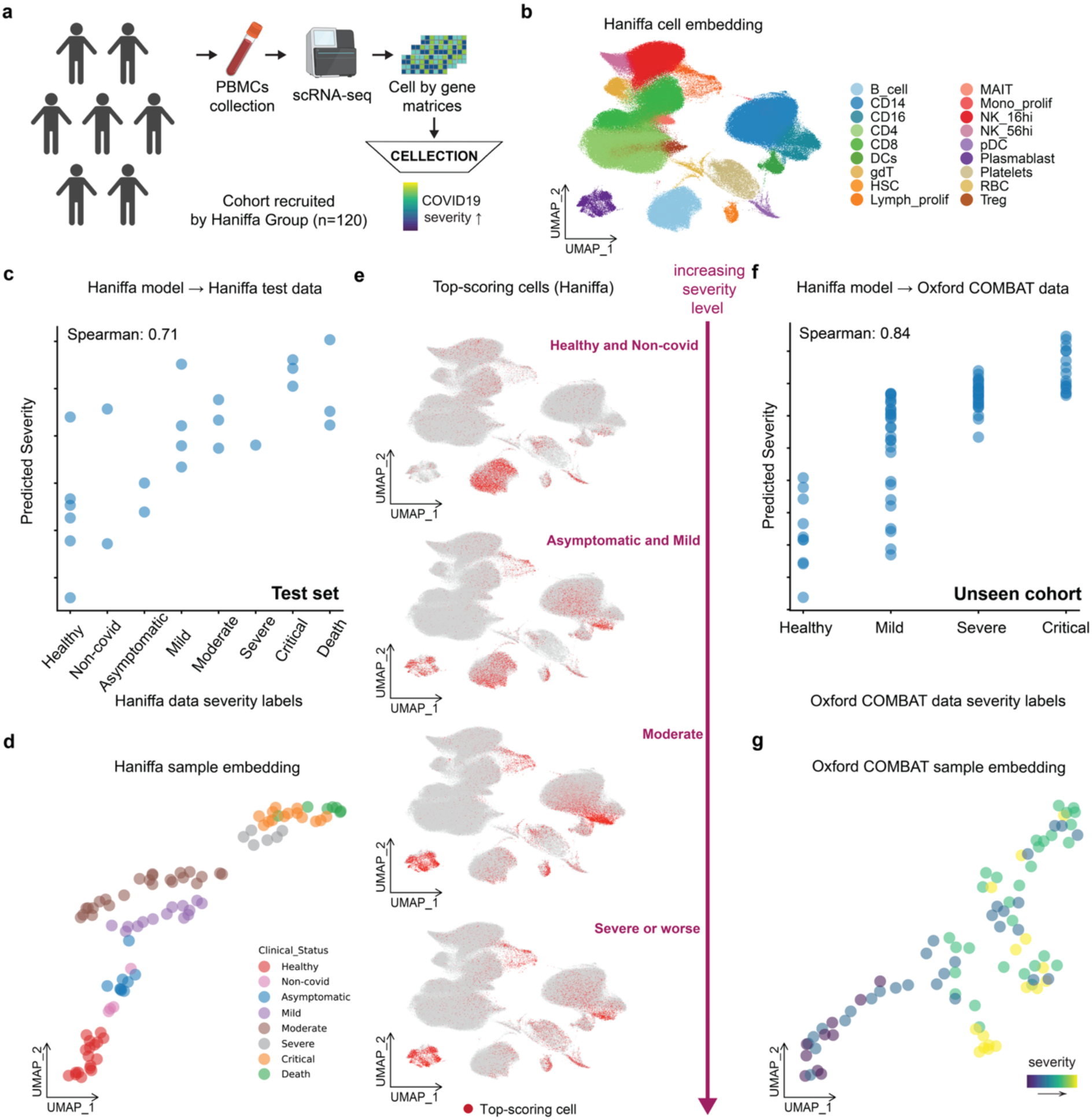
Sample embeddings capture COVID-19 severity progression. (**a**) Overview of the prediction experiment design. PBMCs were collected from COVID-19 patients in the cohort recruited by Haniffa Group and profiled using scRNA-seq^30^. Samples were labeled with ordinal severity levels and used to train CELLECTION for severity prediction. (**b**) UMAP of CELLECTION cell-level embeddings from the Haniffa dataset, colored by original cell type annotations. (**c**) CELLECTION performance on test samples from the Haniffa cohort, showing a Spearman correlation of 0.71 between predicted and true severity. (**d**) Sample-level embeddings for Haniffa data, colored by annotated disease severity. (**e**) Cell-level UMAP from Haniffa data, with the top 10% highest-scoring cells per sample highlighted in red, showing cell prioritization patterns across severity conditions. (**f**) Transfer learning performance on an independent dataset (Oxford COMBAT)^29^, using the model trained on the Haniffa data. The predicted severity correlates with the ground truth (Spearman = 0.84). (**g**) Sample-level embeddings for the Oxford COMBAT data, colored by severity gradient.

We next assessed generalizability through transfer learning. The Haniffa-trained CELLECTION model was evaluated on the Oxford COMBAT cohort^29^, an independent dataset collected and processed separately from the Haniffa cohort and unseen during training. Despite the domain shift, CELLECTION preserved ordinal structure in its predictions (Spearman = 0.84) (**Figure.3f**) and embedded patient samples along a comparable severity trajectory (**Figure.3g**). These results suggest that CELLECTION captures common underlying and biologically meaningful cellular patterns associated with disease severity, rather than dataset- or cohort-specific artifacts. Importantly, the attention mechanism provided mechanistic insight into which cells were driving the model’s generalized predictions on this unseen Oxford COMBAT cohort. High-attention cells were consistently enriched in immune populations including monocytes and plasmablasts, known to be associated with COVID-19 severity (**Figure.S3c**). These findings are consistent with our results from the Haniffa cohort and align with previous studies^29,30,46–49^, which demonstrated that CELLECTION prioritized major cellular programs along the severity spectrum, mirroring established immunopathological hallmarks of COVID-19.

### Identification of analogous cell types and developmental stages between neural organoids and fetal brains

Building on CELLECTION’s ability to resolve phenotypic gradients in disease progression, we next explored its application in the context of developmental biology. A central goal in developmental biology is to understand how cell types emerge, transition, and organize during development. The brain represents a compelling system for this inquiry, as it undergoes extensive spatial and temporal coordination, with transient progenitor states giving rise to highly specialized neurons and glial populations^50–53^. However, direct *in vivo* measurement of human brain development is constrained by ethical, technical, and logistical limitations^54^. To address this, human neural organoids have emerged as a valuable model system, enabling researchers to observe developmental processes over time in a controlled setting^55,56^.

Despite their promise, organoids often diverge from primary tissue in cell composition, lineage balance, and maturation temporal rates^57,58^. Standard analytical approaches for assessing organoid fidelity rely on clustering cells and comparing cell-type distributions across systems^32,59^. These methods require consistent cell type annotations, but different investigators may apply different rules or use different gene markers to identify cell populations or use different naming conventions^60,61^, or two different studies might sequence at different depths and thus differences between systems can emerge for technical or biological reasons. As a result, it can be challenging to translate findings from specific organoid developmental stages to *in vivo* brain development stages, or vice versa.

Given how poorly nuclei from organoids and developmental brain samples can be integrated and directly mapped to one another (**Figure.4b**), we asked whether CELLECTION ordinal analysis could help map organoids to fetal brain samples, both at the single cell and sample level, in order to uncover shared developmental axes and critical cellular programs. We trained two separate CELLECTION models on single-cell RNA-seq profiles from the Human Developmental Brain Cell Atlas^31^ (HDBCA; n = 340 samples) and the Human Neural Organoid Cell Atlas^32^ (HNOCA; n = 282 samples), each comprising over 1.6 million cells (**Figure.4a**). Using developmental age as the prediction target (organoid time in days or fetal age in post-conception weeks), CELLECTION accurately recovered sample-level trajectories within each system (Spearman = 0.96 for HDBCA, and Spearman = 0.99 for HNOCA on the validation set; **Figure.S4a, b**). These results confirm that transcriptomic signals of maturation are robustly encoded in population-level cell distributions.

**Figure. 4.**
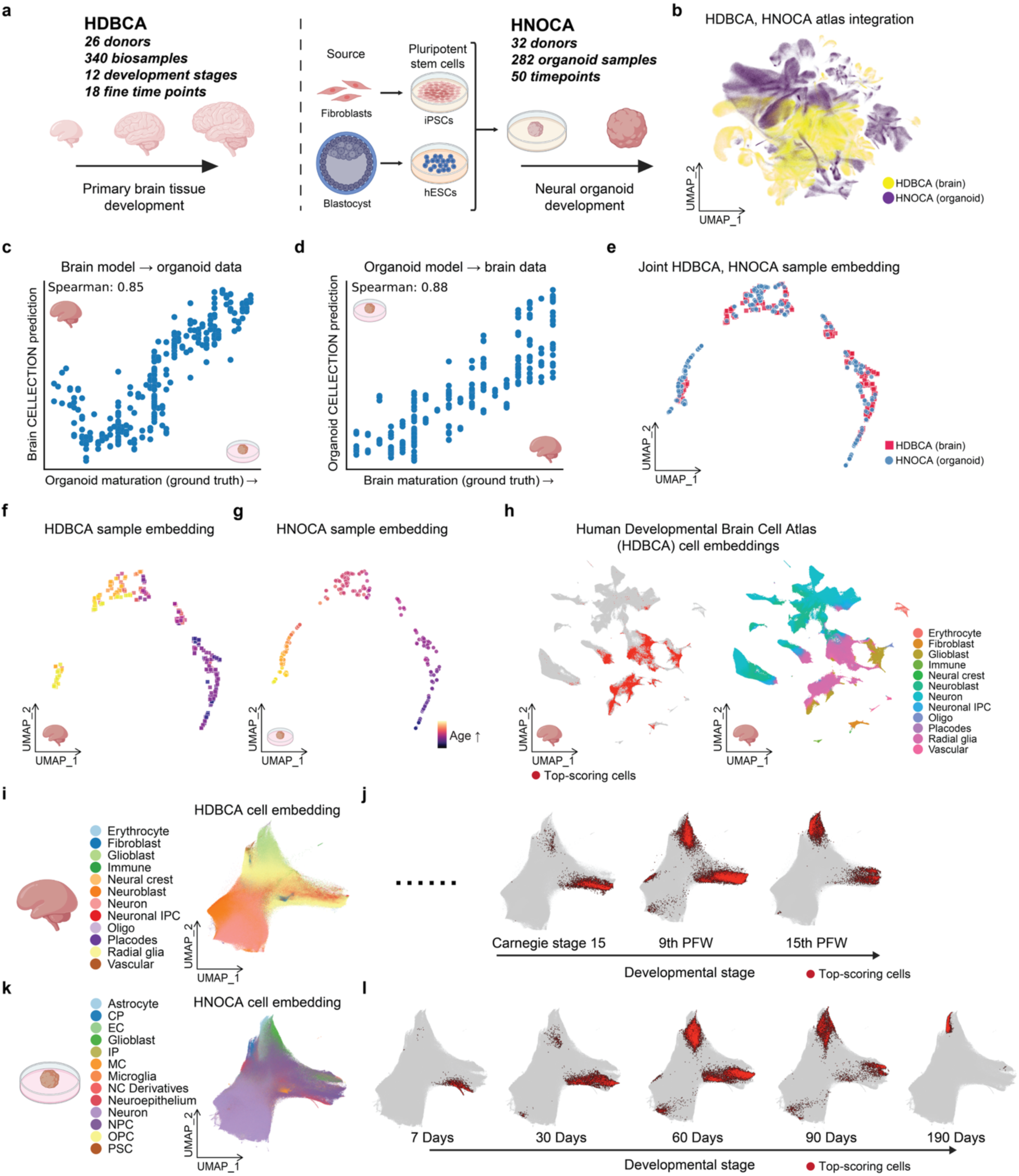
Identification of analogous developmental stages and cell populations between neural organoids and fetal brains. (**a**) Schematic overview of the two atlases used in this experiment: the Human Developmental Brain Cell Atlas (HDBCA)^31^ comprising fetal brain samples across 12 developmental stages, and the Human Neural Organoid Cell Atlas (HNOCA)^32^ comprising organoid samples across 50 differentiation time points. (**b**) Cell-level UMAP shows poor per-cell transcriptome overlap between neural organoids and fetal brain samples. (**c-d**) Cross-system prediction performance using ordinal regression. Models were trained on one system (e.g., organoid) and applied to the other (e.g., fetal brain) to test generalization, with Spearman correlations of 0.85 (primary brain model tested on organoid) and 0.88 (organoid model tested on primary brain), respectively. (**e**) CELLECTION sample-level embedding of HDBCA and HNOCA samples, showing convergence of developmental trajectories across the two systems. (**f-g**) Sample-level embeddings for HDBCA (**f**) and HNOCA (**g**) individually, colored by annotated developmental or maturation stages. (**h**) Cell-level UMAPs from the HDBCA. Left: cells with the highest 10% of scores are highlighted in red, indicating their contribution to developmental stage prediction. Right: the same embedding but colored by original cell type annotations from the HDBCA. (**i-l**) Summary of analogous cell populations identified across developmental stages of organoids and fetal brains: (**i**) cell-level embedding of the HDBCA learned by CELLECTION; (**j**) representative developmental stages from fetal brain with high-scoring cells for each stage in red; (**k**) cell-level embedding of the HNOCA learned by CELLECTION; (**l**) representative timepoints during organoid maturation with high-scoring cells for each timepoint in red.

To evaluate cross-system generalization, we applied the CELLECTION model trained on one atlas to predict developmental age in the other atlas. CELLECTION maintained strong performance in both directions (brain model predicts organoid age: Spearman = 0.85; organoid model predicts brain age: Spearman = 0.88) (**Figure.4c, d**), suggesting that CELLECTION captures common underlying transcriptomic programs shared across biological systems and technical conditions. Notably, the sample-level embeddings learned by CELLECTION revealed a smooth and convergent developmental axis, with organoid and brain samples aligning according to their respective stages of maturation (**Figure.4e-g**).

In order to identify the cellular programs underlying this developmental alignment of organoids and brains, we used the CELLECTION model trained on HNOCA data, which provides broader temporal coverage, to identify high-scoring cells that contributed most to stage prediction in both datasets. For early developmental stages, radial glia drove predictions for primary fetal brain samples (**Figure.4h, i, j** and **Figure.S4**) while neural progenitor cells (NPCs) and neuroepithelial cells drove predictions in organoids (**Figure.4k, l**). This is consistent with how radial glial cells function similarly to NPCs by undergoing asymmetric divisions to produce neurons directly or indirectly through intermediate progenitor cells^62–65^. As development progressed, CELLECTION attention shifted to glioblasts in both systems (**Figure.4h-l** and **Figure.S4c**). Due to the larger sample size and resulting temporal coverage of the organoid dataset, CELLECTION also identified mature glial cell states emerging beyond day 90 for the organoids (**Figure.4l**), reflecting later fate commitments not captured in the fetal brain data due to its limited sample size and temporal coverage. These dynamic attention patterns were learned without access to any cell type labels or clustering, demonstrating CELLECTION’s capacity to uncover conserved developmental hierarchies in a data-driven manner. Importantly, the model learned a unified embedding space in which samples from both systems aligned by maturation stage (**Figure.4e**) and similarly for cells (**Figure.4i, k**), even though the original studies used distinct annotation protocols and naming conventions, and the cells from the original studies cannot be reliably integrated (**Figure.4b**). This cross-system alignment enabled the discovery of analogous cell populations across organoids and fetal tissue, without requiring extensive integration, batch correction, or manual curation.

To further demonstrate the utility of CELLECTION in organoid systems, we applied the model to predict both disease status and donor cell line origin using organoid-derived single-cell data. In the HNOCA disease dataset, which includes 64 samples from three neurodevelopmental disorders (ALS/FTD, ASD, Glioblastoma) and healthy controls, CELLECTION accurately classified disease conditions with high performance (average AUROC = 1.00, Macro F1 = 0.98; **Figure.S5a**). Moreover, in an independent dataset from Paulsen et al.^66^, the model was able to distinguish organoids by their donor cell line of origin, achieving strong separation in the embedding space and robust classification accuracy (AUROC = 1.00, Macro F1 = 0.89). In addition to donor identity, the model also captured temporal variation in the Paulson dataset, by separating time points (1 month, 3 months, 6 months) within each donor line (**Figure.S5b**). These results highlight CELLECTION’s flexibility in modeling not only developmental and disease-related variation, but also technical and experimental sources of heterogeneity directly from organoid-based single-cell data.

### CELLECTION uncovers novel disease-associated microglia signatures in Alzheimer’s disease

We also applied CELLECTION to a neurodegenerative context to investigate how distinct glial cell states contribute to Alzheimer’s disease. We chose Alzheimer’s disease because of evidence suggesting the underlying cellular drivers of pathology are heterogeneous across individuals, despite convergence of their clinical disease phenotype^5–8^. This phenotypic convergence, where multiple cellular programs can give rise to similar clinical outcomes, represents a major challenge for traditional differential expression and cell type proportion analysis, which typically assumes homogeneity within groups. We hypothesized that through analysis of high scoring cells identified by CELLECTION, we could detect and explain AD patient heterogeneity through systematic differences in the cell types of the high scoring cells.

Glial cell populations are widely recognized as key players in neurodegeneration through their roles in inflammation, myelination, and metabolic support^67–73^. This includes microglia, oligodendrocytes, oligodendrocyte precursor cells (OPCs), and astrocytes (**Figure.5b**). These cell types are inherently heterogeneous, prompting the question of whether specific subtypes of glial cells are responsible for AD pathology. To investigate this, we trained CELLECTION on the Seattle Alzheimer’s Disease Brain Cell Atlas (SEA-AD) dataset^25^, which includes snRNA-seq profiles from the middle temporal gyrus (MTG) of 87 human postmortem brains spanning both AD cases and controls (**Figure.5a**). We trained it to predict Alzheimer’s case-control status using only glial cell populations from each individual sample. In this initial analysis using all glial cell populations together, CELLECTION achieved strong predictive performance on the validation set (AUROC = 0.87, Macro F1 = 0.82), without requiring any predefined cell type labels. Its performance is significantly higher that of a model using cell type proportions alone (AUROC = 0.70, Macro F1 = 0.49), suggesting for SEA-AD glial populations, CELLECTION is capturing more than just differences in sample composition.

**Figure. 5.**
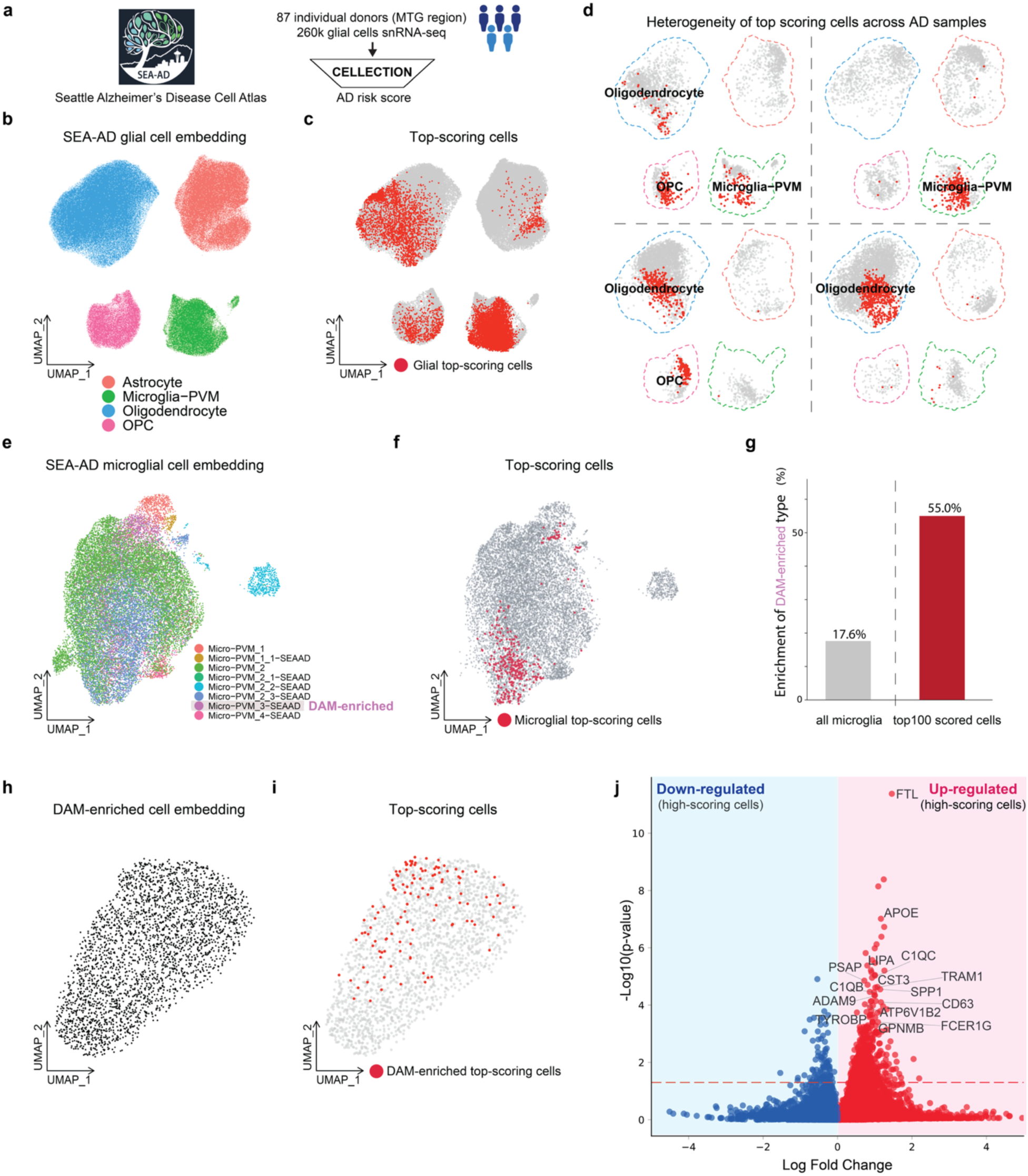
CELLECTION identifies novel microglial cell states associated with Alzheimer’s disease. (**a**) Overview of the subset of the Seattle Alzheimer’s Disease Brain Cell Atlas (SEA-AD) used to train CELLECTION, consisting of over 260k glial cells from 87 individual donors. (**b**) Cell-level UMAP of all glial cells, colored by annotated glial subtypes: astrocytes, microglia, oligodendrocytes, and oligodendrocyte progenitor cells (OPCs). (**c**) Cell-level UMAP of glial cells from AD donors, with the top 10% high-scoring (AD-predictive) cells highlighted in red for each individual. (**d**) Donor-specific UMAPs of high-scoring cells for four donors, showing heterogeneity in prioritized glial subpopulations across AD patients. (**e**) Cell-level UMAP of the microglia subset of cells from all donors of SEA-AD, colored by microglial subtype annotations from the original SEA-AD study. The microglial subtype Micro-PVM_3-SEAAD contains disease-associated microglia (DAM), and is named the DAM-enriched cell state here. (**f**) Same as (e), but the top 10% highest-scoring cells predictive of AD status are colored in red. (**g**) Bar plot showing the proportion of DAM-enriched cells among all microglia (17%) is much less compared to among highest scoring cells identified by CELLECTION (55%). (**h**) Cell-level UMAP of the DAM-enriched microglial subset of SEA-AD, shown without further sub-annotation as none are provided in SEA-AD. (**i**) Same as (h), but the top 10% highest-scoring cells predictive of AD status are colored in red. The non-uniformly distributed red cells suggest the presence of a refined DAM-like subpopulation. (**j**) Volcano plot of differentially expressed genes (DEG) in the high-scoring DAM-like subpopulation identified by CELLECTION, relative to other DAM-enriched cells. Known AD markers such as *APOE*, *SPP1*, *FTL*, and *C1QB/C* are significantly upregulated.

To investigate which glial cell populations overall were responsible for this high prediction accuracy, we identified the high-scoring cells across all AD cases and found each of the four glial populations made contributions to phenotype predictions for the AD cases as a group (**Figure.5b, c**). To investigate the extent of patient-level heterogeneity in the AD cases, for each individual AD brain sample we identified the high-scoring cells responsible for AD prediction for that case only, and identified the major cell types of only the high-scoring cells for each sample. In many patients, microglia were the dominant contributors (**Figure.S6a, b**), while in others, astrocytes, oligodendrocytes, OPCs, or combinations thereof played the primary role (**Figure.5d**). This variation suggests that AD may not be driven by a single glial program, but rather by heterogeneous cellular mechanisms across individuals, which reflects underlying transcriptional heterogeneity in disease manifestation. Overall, the majority of the AD patients (79%) were predicted as AD primarily due to their microglia cell states, suggesting at least within the SEA-AD dataset, changes to microglia states are strong molecular signatures of AD.

To more precisely determine what changes in microglia are associated with AD status, we trained CELLECTION on the SEA-AD dataset again, however this time using only microglia cells from each patient (**Figure.5e**) instead of all glial cells as in the previous analysis. By doing so, we are effectively conditioning our analysis on the observation that microglia are predictive of case-control status, and now asking to what extent variation within microglial populations specifically explains AD case-control status. Even using microglia cells alone, CELLECTION maintained strong predictive performance on the validation set (AUROC = 0.77, Macro F1 = 0.75) compared to using cell subtype proportions (AUROC = 0.62, Macro F1 = 0.52).

Using the microglia-trained CELLECTION, we identified high-scoring cells driving the high AD prediction accuracy (**Figure.5e, f**). The top-scoring cells were enriched in the microglial subtype annotated as the “DAM-enriched” (disease-associated microglia) cluster previously described in the SEA-AD study (specifically, the Microglia-PVM_cluster 3_SEAAD cluster, which was reported to contain DAMs alongside related reactive states)^25^ (**Figure.5e, f**). Although the DAM-enriched cluster accounted for only 17% of the total sequenced microglial population, these cells represented more than 55% of the top-scoring, AD-driving microglia identified by CELLECTION (**Figure.5g** and **Figure.S6c**). This strong enrichment agrees with prior findings from SEA-AD and highlights CELLECTION’s ability to recover established disease-relevant states through attention-based prioritization, without relying on traditional analysis pipelines that require extensive preprocessing, clustering, manual curation, and re-clustering.

In the original SEA-AD study, the identification of the DAM-enriched cluster represented the finest resolution used to characterize disease-associated microglia. As a clustering- and annotation-free method, CELLECTION potentially can identify specific cell states even amongst the disease-associated microglia that more precisely pinpoint the disease signature. To do so, we repeated CELLECTION training on the SEA-AD dataset again, but further restricted the model input to only DAM-enriched microglia cells from each patient (**Figure.5h**). Surprisingly, the model retained strong predictive power on the validation set (AUROC = 0.77, Macro F1 = 0.67) which was more predictive than or cell count in this case (AUROC = 0.61, Macro F1 = 0.48), indicating that even within DAM-enriched microglia cells, more precise and functionally relevant subpopulations can be robustly identified by CELLECTION (**Figure.5i**). A differential expression analysis of the top-scoring subset of the microglial subcluster input revealed enrichment for well-established DAM markers^74,75^, including *APOE*, *SPP1*, *FTL*, and members of the *C1Q* gene family (**Figure.5j**), suggesting that CELLECTION can uncover finer-grained, precise disease-predictive cell states beyond those captured by clustering-based approaches. Our results here thus suggest the existence of a novel DAM cell signature even more specific than that reported in the original SEA-AD study.

To test whether these disease-predictive microglial signatures could generalize beyond SEA-AD, we applied the microglia-trained CELLECTION model to the ROSMAP dataset (comprising 930,739 nuclei from 349 individual subjects) directly, without fine-tuning on the ROSMAP cohort. We only tested the microglia-trained CELLECTION but not the DAM-enriched CELLECTION because the cell annotations of the ROSMAP dataset only resolve cells up to a microglial cell state. Notably, despite differences in brain region and disease cohort between SEA-AD and ROSMAP, CELLECTION was predictive on the ROSMAP microglial data (AUROC = 0.61, Macro F1 = 0.58), outperforming baseline models that relied on cell counts, subtype proportions, or their combination (**Figure.S7a**). To investigate what cell states were prioritized in this held-out dataset, we visualized the top-scoring microglia from both AD and normal control samples using sciLaMA-based UMAP embeddings. Here, the sciLaMA model was trained jointly on SEA-AD and ROSMAP microglial data to learn a shared low-dimensional embedding space, which enabled more efficient transfer learning by skipping the need to retrain CELLECTION’s InstanceEncoder. In the resulting embedding, top-scoring cells from AD and normal samples formed largely non-overlapping clusters (**Figure.S7b**), suggesting that CELLECTION effectively distinguishes distinct cellular programs associated with AD status. We further characterized these prioritized cells using differential gene expression (DEG) analysis. Among the top-scoring cells from AD samples, we observed significant upregulation of canonical DAM markers, including *APOE*, *ACSL1*, while top-scoring cells from normal controls showed downregulation of DAM markers such as *C1QB* and *C1QC* (**Figure.S7c-e**). These findings reinforce that CELLECTION not only generalizes well across cohorts but also prioritizes molecularly distinct, disease-relevant subpopulations in a clustering- and annotation-free manner.

### CELLECTION generalizes to phenotype prediction at evolutionary scales

Finally, we wanted to test the extent to which CELLECTION could be applied to other emergent phenotype prediction problems. To that end, we applied it to model how variation in genome content could drive phenotypic variation at evolutionary scales. Over the past few years, a number of large language models have been developed to model variation in whole genome sequence^76–80^. While they are often trained in an unsupervised manner, they have been fine-tuned to address supervised prediction tasks such as promoter and motif identification, splice site discovery, and genetic variant effect estimation^81^. However, these LLMs model the genome at nucleotide-level resolution, which can be challenging for long genomes such as the human genome. Here, we hypothesized that we could treat a genome as an unordered collection of genes, with which we associate organism-level phenotypes. To test CELLECTION under this scenario, we needed to acquire training data consisting of many assembled, annotated genomes with consistently measured phenotypes across those same organisms. Furthermore, we desired to predict phenotypes for complex genomes such as vertebrates. The Bird 10,000 Genomes (B10K) Project^82,83^ has assembled and annotated 265 avian genomes, for which 201 genomes also have morphological measurements from the AVONET database^84^. More specifically, we used the hand-wing index (HWI)^85^, a morphological trait reflecting avian flight efficiency, as the target phenotype, and treated each bird species as a bag of gene-level embeddings derived from the protein language model ESM2^86^. CELLECTION performed well at these phenotype predictions as well, achieving a Spearman correlation of 0.89 on the training set and 0.57 on the testing set (**Figure.6**). These results demonstrate that CELLECTION can generalize from cellular to organismal scales, providing a unified framework for linking molecular-level features to high-level phenotypes across biological hierarchies.

**Figure. 6.**
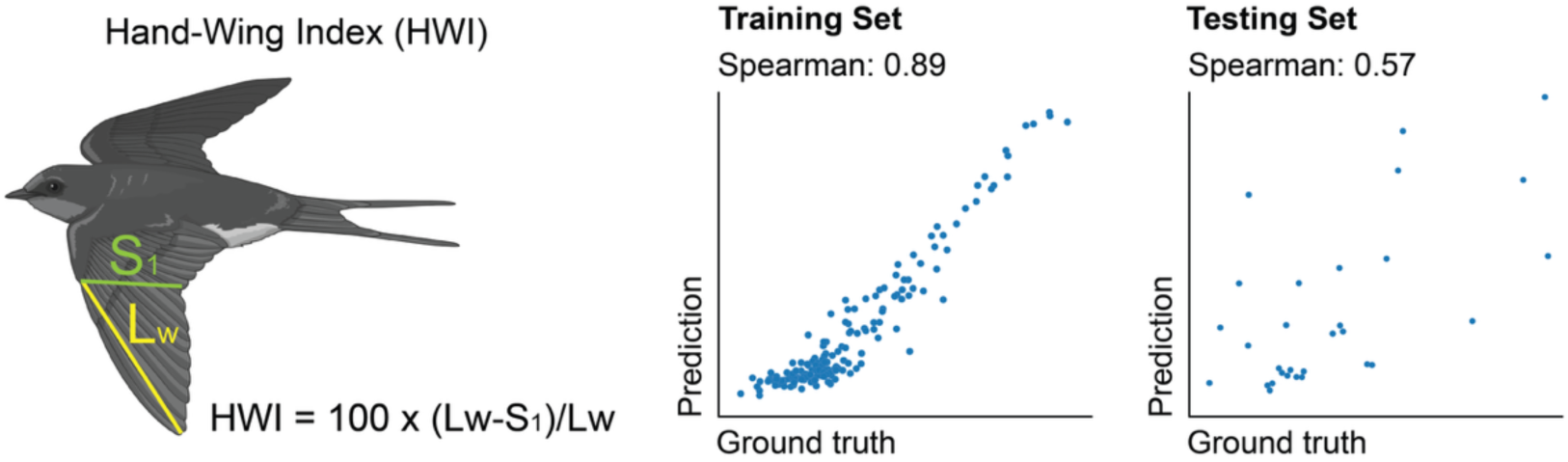
Predicting organism-level phenotypes. The CELLECTION architecture can scale beyond single-cell data to genome-level inference by treating genes as instances of a whole genome. Shown here is an application to predict hand-wing index (HWI), a quantitative trait related to avian flight efficiency. Scatter plots show model performance on the training set (left, Spearman = 0.89) and testing set (right, Spearman = 0.57) from a set of 265 avian genomes, demonstrating the model’s capacity to predict organismal phenotypes from genomic content.

## Discussion

A central challenge in biology is understanding how complex phenotypes emerge from diverse and heterogeneous cellular populations. CELLECTION offers a powerful framework to address this by directly linking single-cell profiles to sample-level outcomes, without relying on predefined cell types, manual clustering, or extensive preprocessing. Importantly, it enables biological discovery by identifying the specific subpopulations that drive phenotypic variation. Our results demonstrated that CELLECTION can identify analogous cell populations and developmental time points between human neural organoids and primary fetal brain tissue, despite the lack of transcriptional similarities of individual cells across the two systems. In the context of neurodegenerative disease, CELLECTION identified a refined disease-associated microglial (DAM) cell signature in Alzheimer’s disease that is more precise than current disease-associated microglial signatures, highlighting its ability to reveal novel, more precise and functionally relevant cellular states. By learning from cellular profiles and assigning importance scores to individual cells, CELLECTION provides interpretable outputs that guide hypothesis generation and offer mechanistic insight into disease and development.

One of the persistent challenges in large-scale single-cell studies is the lack of standardized cell type annotations across datasets, which often differ in resolution and clustering strategies, or rely on inconsistent naming protocols. In the Human Neural Organoid Cell Atlas (HNOCA) and the Human Developmental Brain Cell Atlas (HDBCA)^31,32^, for example, cells representing similar developmental lineages are assigned distinct, non-overlapping labels in the original studies. This stems from the fact that these datasets originate from different biological systems and studies, and therefore underwent different annotation protocols. As we showed in **Figure.4b**, even after integration it can be difficult to identify matching cell types across different annotations. The Human Cell Atlas (HCA) consortium has highlighted that definitions of “cell type” and “cell state” remain imprecise, and that current annotations often lack the granularity needed for consistent cross-dataset comparison^87^. CELLECTION bypasses these annotation bottlenecks at the cell-instance level by making predictions without cell type labels. Instead, it learns to identify shared cellular programs directly from the cellular profiles, using sample-level phenotypes as weak supervision. This makes it particularly valuable for analyzing samples or tissues that are poorly studied, aligning datasets with sparse or inconsistent annotations, comparing samples across species, or identifying cell populations characteristic of specific phenotypes without the need to construct a unified reference cell atlas.

Several methods have previously aimed to extract phenotype-relevant signals from single-cell data^38–40^, but CELLECTION addresses key limitations that constrain their performance and interpretability as we demonstrated in our benchmarking results. For example, CloudPred fails to capture the variable importance of individual cells; CELLECTION instead employs an attention mechanism to weight cells by their contribution to phenotype, enabling easy identification of cell states associated with different phenotypes. ProtCell4P enhances interpretability through prototype learning but relies on predefined cell type annotations and a fixed number of prototypes, limiting its flexibility and application to datasets for which cell type annotations are available and at high resolution. In contrast, CELLECTION requires no such labels and adaptively identifies relevant subpopulations directly from raw data. DeepSet all instances (cells) equally through sum pooling, which can obscure rare yet critical cell states. CELLECTION builds on this foundation with a more expressive cell embedding architecture and attention-based aggregation, allowing it to highlight informative cells without diluting subtle signals.

As we demonstrate in **Figure.6** with the prediction of the hand-wing index (HWI) for avian genomes, CELLECTION is readily applied to make emergent phenotype predictions outside the context of single cell biology. In the case of HWI, the input instances were genes encoded in a specific avian genome, instead of cells sequenced from a sample. In other words, CELLECTION generalizes to genome-scale data by treating genes as instances to predict organism-level traits, and could be readily used to study the evolution of traits across phylogenies.

A common framework for linking genetic variation to phenotypic variation is the polygenic risk score (PRS), which is typically calculated as the weighted sum of risk alleles in an individual’s genome, where alleles are weights by their estimated effect size. The PRS is therefore a statistic that represents the relative genetic risk of an individual for a disease. Through the use of its encoder, attention, aggregation and fully connected network, CELLECTION could be viewed as a nonlinear form of PRS for complex phenotypes. We therefore envision CELLECTION being applied in the context of genetics as a sophisticated PRS, where variants are encoded by genomic features such as epigenomic or transcription factor binding site profiles, or coding features (in the case of protein-coding variants) in order to compute PRSes that are context-aware.

CELLECTION is a highly extensible framework in that it can readily incorporate pretrained instance-level embeddings from multimodal or foundation models in place of the its own instance encoder in order to leverage multimodal knowledge and external biological priors^36,88^ (based on the training data of the foundation model) to support discovery.

In our experiments, we leveraged CELLECTION to establish direct analogies, both at the individual cell type level and developmental stage, between human neural organoids and fetal brains. CELLECTION therefore offers strategies for evaluating organoid fidelity, identifying when and how organoids diverge from *in vivo* brain development, and probing the cellular programs that govern early human brain maturation. Furthermore, it holds particular value in disease modeling and therapeutic screening. For instance, researchers studying early-stage neurodevelopmental disorders can use CELLECTION to pinpoint which organoid timepoints most closely mirror critical *in vivo* windows. This capability can guide the timing of perturbation assays, therapeutic interventions, or phenotype assessments in disease-relevant *in vitro* systems, and can further provide a concrete path toward more precise and actionable experimental design.

## Methods

### The CELLECTION framework

CELLECTION is a deep multiple instance learning (MIL) architecture for predicting phenotypes from variable-length sets of biological instances, such as collections of cells corresponding to multiple individuals in a population-scale single-cell omics datasets, or gene catalogs corresponding to individual genomes in comparative genomics studies (**Figure.1a**). It adapts principles from PointNet^37^ to biological data by treating each sample as an unordered collection of high-dimensional feature vectors, making the framework inherently permutation-invariant. CELLECTION functions in a cell-level annotation- and clustering-free fashion, which means it does not rely on predefined cell types, manual labeling, or external clustering steps. Instead, it processes all instances (cells) directly, preserving single-cell resolution while enabling end-to-end learning of sample-level representations.

Let ***X*** ∈ ℝ^*N*×*G*^ represent a single sample input to CELLECTION, consisting of a collection of *N* instances (e.g. single cells), each of length *G* (number of genes). Initially, each instance (cell) ***X***_*i*_ is first passed through an optional instance encoder to produce a low-dimensional representation 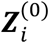, where ***Z***^(0)^ ∈ ℝ^*N*×*F*^:

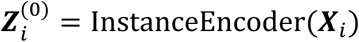

Where InstanceEncoder maps the *G* input features to a low, *F*-dimensional representation through a stack of fully connected (FC) layers with customizable activation functions. It optionally supports layer normalization^89^, batch normalization^90^, and dropout^91^ for regularization. Batch covariates are concatenated to each instance and encoded jointly, allowing the model to correct for batch effects when available. Alternatively, pretrained representations from other frameworks can be used in place of the instance encoder, such as sciLaMA^36^ which leverages multimodal large language models (LLMs), or scPair^88^ which supports single cell multiomics data including 10x Genomics scMultiome and CITE-seq. In the benchmarking experiments, the InstanceEncoder was used for all datasets, using a two-layer architecture with hidden dimensions of 256 and 32 by default. Each layer consisted of a linear transformation followed by layer normalization, ReLU activation, and a dropout rate of 0.1. For the Alzheimer’s disease (AD) experiments, sciLaMA was first trained to learn instance-level low-dimensional representations, where its cell encoder contained two hidden layers of size 1500 and 300, followed by a projection to a 50-dimensional embedding space. After sciLaMA training on the initial scRNA-seq data, we used its encoder in place of the InstanceEncoder in CELLECTION.

After passing individual instances through the optional instance encoder, each instance is then passed through *K* = 2 consecutive Feature Transformation Blocks (**Figure.1a**). The *k*^th^ Feature Transformation Block learns and applies an *F*-dimensional affine transformation ***T***^(*k*)^to the encoded input data from the previous Feature Transformation Block ***Z***^(*k*–**1**)^, to make the data ***Z***^(*k*–**1**)^ more comparable across samples:

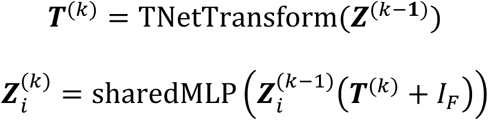

Where *I*_*F*_ is an *F*-dimensional identity matrix. The TNetTransform network applies a series of 1D convolutions using kernels of dimension *F* across the instance axis of ***Z***^(*k*–**1**)^, followed by max pooling and fully connected layers, whose outputs are re-shaped into the transformation matrix ***T***^(*k*)^ ∈ ℝ^*F*×*F*^. ***T***^(*k*)^ is then added to the identity matrix *I*_*F*_ before multiplication with 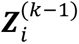 to encourage transformations that are close to the identity matrix (no transformation). The product ***Z***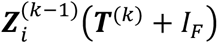 is finally passed through a shared-weight multi-layer perceptron (sharedMLP) before the output of the *k*^th^ Feature Transformation Block 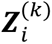 is computed. We stack two of these Feature Transformation Blocks by default in our experiments, to enhance feature alignment across samples.

Following the *K* Feature Transformation Blocks, the processed instances ***Z***^(*K*)^ are then passed through a Weighted Aggregation Block to summarize the entire sample into a single sample-level embedding vector ***s*** ∈ ℝ^*F*^, regardless of the number of input instances. CELLECTION supports several aggregation strategies. The primary and most expressive approach uses attention-based mechanisms, in which CELLECTION computes learnable importance weights (a.k.a. cell scores in single cell data) for each instance. The two main attention formulations are:

(1) standard attention (using tanh activation):

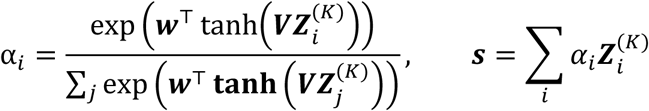
(2) gated attention (combining tanh and sigmoid activations):

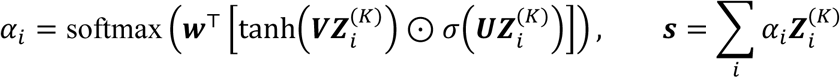

Here, 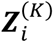 denotes the representation of the *i*-th instance after the final *K*^th^ Feature Transformation Block, and the parameters ***V***, ***U***, ***w*** are learned during training. In our implementation, the attention parameters are configured as follows: ***V*** and ***U*** are weight matrices of shape (64, *d*_01_), and ***w*** is a vector of shape (64, 1), where *d*_01_ corresponds to the dimensionality of the instance representation after the final Feature Transformation Block. In addition to attention, CELLECTION also supports symmetric pooling operations as alternative aggregation methods such as max, mean, and sum pooling. During inference, the model retains interpretable outputs, including attention weights for attention-based aggregation, identifying high-importance instances, or critical indices for max pooling, revealing the most informative instances. The resulting sample-level embedding ***s*** is passed to a sample-level predictor network PhenoMLP, by default a shallow MLP with one hidden layer. This module transforms ***s*** into the final output, such as disease classification, trait regression, or severity ranking, depending on the task.

### CELLECTION data standardization

Each input sample in CELLECTION is represented as ***X*** ∈ ℝ^*N*×*G*^, a tensor where the rows correspond to individual biological instances, such as cells in single cell data or genes in genomic data, and the columns represent their feature dimensions (e.g., gene expression values or pre-defined embeddings). Since different samples may contain varying numbers of instances, CELLECTION uses zero-padding to standardize input shapes during batching. A corresponding binary mask is applied to distinguish real entries from padding, ensuring that padded positions are excluded from attention score calculations and loss evaluation.

### CELLECTION training

The CELLECTION framework is trained using a stochastic gradient descent strategy, the Adam optimizer^92^. Both the model architecture and training procedure are implemented using PyTorch^93^, with model parameters optimized using the Adam optimizer and a default learning rate of 1e-4. By default, training is conducted for up to 200 epochs, with early stopping triggered if the validation loss does not improve for 20 consecutive epochs. The model checkpoint corresponding to the best validation loss is retained for evaluation. A mini-batch size of 5 samples is used for benchmarking experiments, while 15 samples are used in case study analyses. Each fully connected layer in the model comprises a linear transformation, followed by layer normalization, a ReLU activation function, and dropout regularization with a rate of 0.1. All weights are initialized using Xavier normal initialization^94^.

The loss function is selected according to the prediction task: (i) for classification, a softmax activation is used in conjunction with categorical cross-entropy loss, (ii) for continuous regression tasks, a linear output layer is trained using mean squared error (MSE), (iii) for ordinal regression, CELLECTION incorporates a cumulative link model that maps a continuous latent score to discrete ordinal categories using a set of fixed thresholds (cutpoints). These cutpoints are initialized as evenly spaced values and are numerically ordered (e.g., via torch.arange), ensuring that the model respects the ordinal structure of the target variable.

To ensure robustness and generalizability, each experiment is repeated across multiple random seeds and cross-validation folds. By default, we use 10 training seeds for benchmarking experiments and 5 seeds for all other settings, combined with 5-fold cross-validation. Reported metrics reflect the average across these multiple runs.

When applicable, pretrained cell or gene embeddings (e.g., from sciLaMA^36^, ESM2^86^, scPair^88^, and etc.) are used as input features and in those cases, the InstanceEncoder is avoided. These embeddings are held fixed during training unless otherwise stated. Batch effects or donor-specific covariates can be optionally appended as additional features to the input layer to control for known confounding variables.

### CELLECTION ordinal loss function

Let *o*_*i*_ ∈ ℝ be the scalar, linear predictor output of the last layer of PhenoMLP for sample *i*, *y*_*i*_ be the final class label of sample *i*, and let *C* be the number of ordinal classes. We define a set of ordered cutpoints *θ*_1_, *θ*_2_, … , *θ*_*C*–1_ and use the sigmoid function σ(⋅) to construct cumulative probabilities:

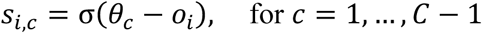

Then, the probability that sample *i* belongs to class *c* is defined as:

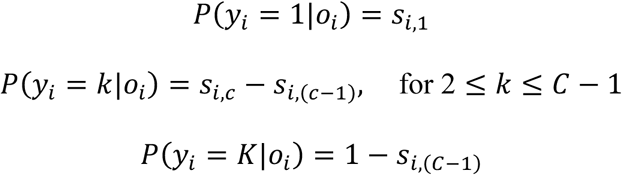

To stabilize training, we clip all probabilities to a minimum value *ε* > 0 to avoid taking the log of zero. The final negative log-likelihood loss is computed as:

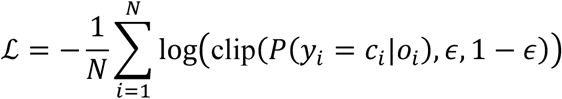

This formulation encourages the model to assign high confidence to the correct ordinal category while respecting the rank structure between neighboring classes.

### Curvature Smoothness (CS) Score

To quantitatively assess how well sample-level embeddings capture an ordered biological trajectory, such as disease severity and developmental stages, we computed a Curvature Smoothness (CS) Score. This metric measures the continuity and geometric smoothness of the trajectory formed by centroids of phenotype classes (e.g., clinical severity levels) in the embedding space.

For each phenotype group, we computed the centroid *C* of its sample-level embeddings. These centroids were then ordered according to their known biological progression, and the first-order differences can be represented as:

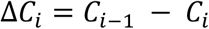

The CS score is derived from the second-order finite differences between consecutive centroids, capturing deviations in curvature along their ordered path.

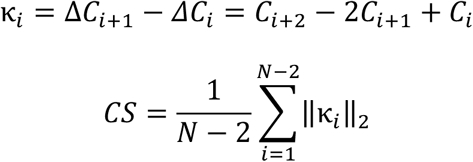

Lower CS scores indicate smoother, more continuous progression in the sample-level embedding space, which is desirable for modeling ordered phenotypes like disease severity and developmental stages.

### Execution of benchmarking methods

We compared CELLECTION to baseline and state-of-the-art models in our benchmarking experiments. These included linear prediction (sklearn.linear_model^95^) using pseudo-bulk averaging, cell type-specific pseudo-bulk averaging, and cell type proportion vectors, as well as the deep learning DeepSet, CloudPred, and ProtCell4P with 4, 8, and 16 prototype configurations. All models were trained and validated using identical data splits for each prediction task to ensure comparability. We implemented benchmarking baselines according to publicly available tutorials and best practices provided by the original authors or repositories (https://github.com/manzilzaheer/DeepSets, https://github.com/bryanhe/CloudPred, https://github.com/Teddy-XiongGZ/ProtoCell4P).

### Performance evaluation overview

Model performance was assessed using task-specific metrics. For classification tasks, we report area under the receiver operating characteristic curve (AUROC) and macro-averaged F1 score. For regression and ordinal prediction tasks, we compute both Pearson and Spearman correlation coefficients between predicted and ground-truth values. All reported results represent the average across multiple training seeds and cross-validation folds to ensure robustness and reduce variance due to stochasticity in model training. Specifically, we used 10 random seeds for benchmarking experiments and 5 seeds for all other experiments, combined with 5-fold cross-validation by default.

### sciLaMA execution to learn cell representations

To generate low-dimensional cell-level embeddings in the AD study, we applied sciLaMA^36^, a representation learning framework that jointly learns embeddings for both cells and genes by integrating prior gene knowledge from multiple pretrained large language models with context-specific single-cell data. Static gene embeddings were derived from three sources: the OpenAI text embedding model applied to NCBI Gene summaries^96^, the scGPT single-cell foundation model^97^, and the ESM-2 protein language model^86^. We used the intersection of the gene sets available in these pretrained models and those present in each single-cell dataset. Following the original optimization procedure described in the sciLaMA study, the resulting cell embeddings were then used as input to CELLECTION for downstream phenotype prediction.

### Organism-level gene embedding generation

Protein-coding gene sequences and gene annotations for each species were obtained from the UniProt Reference-Proteome collection (https://ftp.uniprot.org/pub/databases/uniprot/current_release/knowledgebase/reference_proteomes/Eukaryota/) or the NCBI Datasets command-line tools (CLI). Every sequence was passed through the pretrained ESM-2 transformer (esm2_t6_8M_UR50D, 8 M parameters, available from https://huggingface.co/facebook/esm2_t6_8M_UR50D). The final-layer embedding of each protein served as a fixed-length vector representation of its corresponding gene. Thus, for any species *s*, the genome is represented as an unordered set: 𝓖_*s*_ = {𝒈_*s*,1_, 𝒈_*s*,2_, … , 𝒈_*s*,*ns*_}, where *n*_*s*_ is the number of protein-coding genes in species *s*, and 𝒈_*s*,*i*_ is the ESM-2 gene embedding (*d*=320).

### Data acquisition and processing

#### COVID-19 datasets

Single-cell transcriptomic datasets related to COVID-19 were obtained from multiple sources. Standard preprocessing steps were applied to each dataset, including filtering of low-quality cells and cells lacking clear donor or sample-level labels, normalization, and selection of highly variable genes (HVGs), described below for each individual dataset.

The COVID-19 benchmarking dataset was obtained from Ziegler et al.^26^, comprising 32,588 cells and 32,871 genes across 58 samples. Genes expressed in fewer than five cells were filtered out, after which 3,000 highly variable genes (HVGs) were identified using Scanpy (version 1.10.3) with the parameter flavor=“seurat_v3” parameter. Samples were annotated with one of four disease statuses: normal, COVID-19, respiratory failure, and long COVID-19. For binary classification tasks, we grouped these into two categories: normal and COVID-19.

For COVID-19 severity prediction, we used data from the Haniffa group^30^, which profiled 647,366 cells, 24,737 genes, and 192 antibodies across 143 individuals. Samples with clear clinical annotations, including Healthy, Non-COVID, Asymptomatic, Mild, Severe, Critical, and Death were kept, while those labeled as “NaN” or “LSP” were excluded from the analysis, yielding 564,569 cells across 120 donors. In our study, we focused exclusively on gene expression data for modeling severity. We also incorporated data from the Oxford COVID-19 Multi-Omic Blood Atlas (Oxford COMBAT) consortium^29^. This dataset contains 836,148 cells and 37,298 genes. We excluded samples with fewer than 1,000 profiled cells, retaining 85 high-quality donor samples with statuses including Healthy Volunteer (HV), COVID_HCW_MILD, COVID_MILD, COVID_SEV, and COVID_CRIT.

#### Lupus dataset

We obtained publicly available scRNA-seq data from Perez et al.^27^, comprising 834,096 cells and 32,738 genes. Genes expressed in fewer than five cells were excluded, resulting in a final gene set of 24,205 genes. We then identified 3,000 highly variable genes (HVGs) using Scanpy (version 1.10.3) with the parameter flavor=“seurat_v3”. The dataset includes peripheral blood mononuclear cells (PBMCs) from 169 individuals, including 50 healthy controls and 119 diagnosed with systemic lupus erythematosus (SLE). For ancestry-related analyses, samples were categorized into 106 White and 63 Asian donors based on reported ethnicity.

#### Cardio dataset

We obtained publicly available scRNA-seq data from Chaffin et al.^28^, which profiled 592,689 cells across 36,601 genes from human heart tissue. After filtering out genes expressed in fewer than five cells, 32,151 genes were retained for downstream analysis. We then identified 3,000 highly variable genes (HVGs) using Scanpy (version 1.10.3) with the parameter flavor=“seurat_v3”. The dataset includes samples annotated by both donor-level and biosample-level identifiers. In total, there are 80 biosamples from 42 unique donors. Clinical annotations include three major cardiomyopathy statuses: non-failing (NF) hearts (n = 16), dilated cardiomyopathy (DCM) (n = 11), and hypertrophic cardiomyopathy (HCM) (n = 15).

#### Human Developmental Brain Cell Atlas

The Human Developmental Brain Cell Atlas (HDBCA) dataset^31^ includes single-cell RNA-seq profiles from 1,665,937 fetal brain cells measured across 59,229 genes. After filtering, 1,641,349 high-quality cells from well annotated samples were retained for downstream analysis. Samples were annotated by post-conception week, covering 12 developmental stages and 340 primary brain tissue samples.

#### Human Neural Organoid Cell Atlas

The Human Neural Organoid Cell Atlas (HNOCA) dataset^32^ comprises single-cell RNA-seq profiles from 1,767,674 cells across 36,720 genes, derived from human cerebral organoids spanning a range of developmental time points. After filtering, 1,641,349 high-quality cells from well annotated samples were retained for downstream analysis. Each sample was annotated by its days in culture, covering 50 distinct developmental time points and 282 organoid samples in total. We also obtained the integrated HNOCA disease map from the same study, which includes 12 disease conditions across 89 samples, comprising 409,227 cells and 14,323 genes. For our analysis, we selected the three disease conditions with the largest number of cells, along with control samples, resulting in a subset of 316,108 cells across 64 samples.

#### Paulsen et al., cerebral organoid dataset

We obtained publicly available scRNA-seq data from Paulsen et al.^66^, which includes 99,064 cells and 20,165 genes across 18 individual samples derived from two donor cell lines. The dataset captures organoid development at three harmonized time points: 1 month, 3 months, and 6 months.

#### Seattle Alzheimer’s Disease Brain Cell Atlas (SEA-AD) snRNA-seq dataset

We obtained single-nucleus RNA sequencing (snRNA-seq) data from the Seattle Alzheimer’s Disease Brain Cell Atlas (SEA-AD), specifically the middle temporal gyrus (MTG) region, as published by Gabitto et al.^25^. The glial population subset of the data contains 260,066 nuclei profiled across 33,222 genes from postmortem human brain samples after removing multiomic profiled cells. Clinical metadata includes samples from 87 unique donors, consisting of 42 individuals diagnosed with AD and 45 healthy controls. These were used to train and evaluate models for AD case-control classification, as well as for identifying cell states associated with disease status.

#### Religious Order Study or the Rush Memory and Aging Project snRNA-seq dataset

We obtained single-nucleus RNA sequencing (snRNA-seq) data from the Religious Order Study (ROS) or the Rush Memory and Aging Project (MAP) (together, ROSMAP) from Mathys et al.^6^. The glial population subset of the data contains 930,739 nuclei profiled across 29,268 genes from six different brain regions. Clinical metadata includes samples from 349 unique subjects, consisting of 196 subjects diagnosed with AD and 153 healthy controls. To evaluate generalizability, we applied a pretrained SEA-AD CELLECTION model to this dataset by uniformly randomly downsampling the number of nuclei per subject to match the minimum number of glial cells across all individuals, thereby controlling for cell count effects. We also trained a linear model on subtype proportions derived from the same downsampled data to serve as a baseline for benchmarking.

#### Hand-Wing Index phenotype dataset of Avian genomes

Genome assemblies for 265 avian species were obtained as part of the Bird 10,000 Genomes (B10K) project (BioProject PRJNA545868: https://www.ncbi.nlm.nih.gov/bioproject/PRJNA545868), which includes high-quality annotated genomes spanning all major avian clades. This dataset was originally compiled by Feng et al.^98^ and later expanded by Stiller et al.^99^, Hand-wing index (HWI) values for 201 overlapping avian species were obtained from the AVONET database compiled by Tobias et al^84^. HWI is a dimensionless morphological metric that quantifies wing pointedness and is widely used as a proxy for flight efficiency and dispersal ability. It is calculated using the following formula from Sheard et al.^85^:

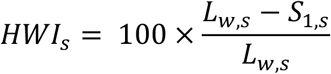

where *L*_w,*s*_ is the wing length (from carpal joint to longest primary) and *S*_1,*s*_ is the length to the first secondary feather, for species *s*. For each species, the HWI value *HWI*_*s*_was used as the regression target variable. The corresponding gene embedding cloud 𝓖_*s*_ was supplied to CELLECTION in regression mode to predict *HWI*_*s*_.

#### Statistics and reproducibility

All statistical calculations were implemented in Python (v3.10.15; https://www.python.org). The detailed statistical tests are indicated in corresponding Figure legends/captions where applicable. No statistical method was used to predetermine sample size. The experiments were not randomized. This study does not involve group allocation that requires blinding.

## Data availability

There is no data generated from this study, and the following publicly available datasets were analyzed in this study:

COVID-19 datasets were obtained from three primary sources. The benchmarking dataset from Ziegler et al. is available via the Broad Institute’s Single Cell Portal:

https://singlecell.broadinstitute.org/single_cell/study/SCP1289. The dataset from Stephenson et al. (Haniffa group) is available through the COVID-19 Cell Atlas (https://covid19cellatlas.org) and can be downloaded from ArrayExpress under accession number E-MTAB-10026. The Oxford COVID-19 Multi-Omic Blood Atlas (Oxford COMBAT) dataset was accessed via the Chan Zuckerberg Initiative (CZI) CELLxGENE Portal at https://cellxgene.cziscience.com/collections/8f126edf-5405-4731-8374-b5ce11f53e82. The lupus dataset was obtained from Perez et al., publicly available through GEO under accession number GSE137029 and the Human Cell Atlas Data Portal at https://data.humancellatlas.org/explore/projects/cc95ff89-2e68-4a08-a234-480eca21ce79. Processed data are available via UCSF Box at https://ucsf.app.box.com/s/tds2gotok3lyeanlrt13prj40am5w720. The cardiomyopathy dataset from Chaffin et al. is available through the Broad Institute’s Single Cell Portal: https://singlecell.broadinstitute.org/single_cell/study/SCP1303. The Human Neural Organoid Cell Atlas (HNOCA) was accessed from the CZI CELLxGENE Portal at https://cellxgene.cziscience.com/collections/de379e5f-52d0-498c-9801-0f850823c847, and the Human Developmental Brain Cell Atlas (HDBCA) was retrieved from https://cellxgene.cziscience.com/collections/4d8fed08-2d6d-4692-b5ea-464f1d072077. The organoid dataset from Paulsen et al. is available via the Broad Institute’s Single Cell Portal: https://singlecell.broadinstitute.org/single_cell/study/SCP1129/. The Seattle Alzheimer’s Disease Brain Cell Atlas (SEA-AD) dataset was accessed via the CZI CELLxGENE Portal at https://cellxgene.cziscience.com/collections/1ca90a2d-2943-483d-b678-b809bf464c30. Only glial cell subsets were used in this study. The Religious Order Study (ROS) or the Rush Memory and Aging Project (MAP) (together, ROSMAP) data were accessed via the AD Knowledge Portal (https://adknowledgeportal.synapse.org) under accession ID syn52293433. Only glial cell subsets were used in this study. Protein-coding gene sequences and annotations for organism-level analyses were retrieved from the UniProt Reference Proteome collection (https://ftp.uniprot.org/pub/databases/uniprot/current_release/knowledgebase/reference_proteomes/Eukaryota/) and the NCBI Datasets command-line interface. https://doi.org/10.1093/jmammal/gyz043. Avian genome assemblies were obtained from the Bird 10,000 Genomes (B10K) Project (BioProject PRJNA545868): https://www.ncbi.nlm.nih.gov/bioproject/PRJNA545868. Hand-wing index (HWI) measurements for bird species were obtained from the AVONET database with the HWI formula defined according to Sheard et al.

## Code availability

The CELLECTION Python package can be installed directly from PyPI: https://pypi.org/project/cellection/. We have made the CELLECTION source code publicly available at https://github.com/quon-titative-biology/CELLECTION, together with a tutorial with sample data provided at https://github.com/quon-titative-biology/CELLECTION/tree/main/tutorials.

## Acknowledgements

This work was supported by NSF CAREER award (1846559 to G.Q.). This has been made possible in part by grants from the National Institutes of Health (NIH), including the Office of the Director/National Institute of Mental Health (DP2 MH129987, G.Q.).

The data available in the AD Knowledge Portal would not be possible without the participation of research volunteers and the contribution of data by collaborating researchers.

## Contributions

G.Q. conceived and supervised the project. H.H. designed the CELLECTION framework and benchmarking strategy. H.H. implemented the CELLECTION prototype, performed all benchmarking and validation analyses, and led manuscript writing. G.Q. and H.H. co-wrote and revised the manuscript, with assistance from S.S. for preprocessing the HWI data. All figures and illustrations were created by H.H. using Adobe Illustrator.

